# Drug treatment efficiency depends on the initial state of activation in nonlinear pathways

**DOI:** 10.1101/153981

**Authors:** Victoria Doldán-Martelli, David G. Míguez

## Abstract

An accurate prediction of the outcome of a given drug treatment requires quantitative values for all parameters and concentrations involved as well as a detailed characterization of the network of interactions where the target molecule is embedded. Here, we present a high-throughput *in silico* screening of all potential networks of three interacting nodes to study the effect of the initial conditions of the network in the efficiency of drug inhibition. Our study shows that most network topologies can induce multiple dose-response curves, where the treatment has an enhanced, reduced or even no effect depending on the initial conditions. The type of dual response observed depends on how the potential bistable regimes interplay with the inhibition of one of the nodes inside a nonlinear pathway architecture. We propose that this dependence of the strength of the drug on the initial state of activation of the pathway may be affecting the outcome and the reproducibility of drug studies and clinical trials.

## Introduction

Some of the main potential contributions of Systems Biology to the field of Pharmacology are to help design better drugs^1,2^, to find better targets^3^ or to optimize treatment strategies^4^. To do that, a number of studies focus on the architecture of the biomolecular interaction networks that regulate signal transduction and how they introduce ultrasensitivity, desensitization, adaptation, spatial symmetry breaking and even oscillatory dynamics^5,6^. To identify the source of these effects, large scale signaling networks are often dissected into minimal sets of recurring interaction patterns called network motifs^7^. Many of these motifs are nonlinear, combining positive and negative feedback and feed-forward loops that introduce a rich variety of dynamic responses to a given stimuli.

In the context of protein-protein interaction networks, these loops of regulation are mainly based on interacting kinases and phosphatases. The strength of these interactions can be modulated by small molecules that can cross the plasma membrane^8^ and block the activity of a given kinase in a highly specific manner^9^. Inhibition of a dysfunctional component of a given pathway via small-molecule inhibition has been successfully used to treat several diseases, such as cancer or auto-immune disorders. Nowadays, 31 of these inhibitors are approved by the FDA, while many more are currently undergoing clinical trials^10^. Characterization of inhibitors and its efficiency^11^ and specificity towards all human kinases constitutes a highly active area of research^12–14^. Importantly, since these inhibitors target interactions that are embedded in highly nonlinear biomolecular networks, the response to treatment is often influenced by the architecture of the network. For instance, treatment with the mTOR-inhibitor rapamycin results in reactivation of the Akt pathway due to the attenuation of the negative feedback regulation by mTORC1^15^, also inducing a new steady state with high Akt phosphorylation^16^. In addition, the nonlinear interactions in the MEK/ERK pathway have been shown to induce different modes of response to inhibition^17^, and even bimodal MAP kinase (ERK) phosphorylation responses after inhibition in T-lymphocytes^18^. The same interplay between positive and negative feedbacks induces ERK activity pulses, with a frequency and amplitude that can be modulated by EGFR (epidermal growth factor receptor) and MEK (Mitogen-activated protein kinase kinase) inhibition, respectively^19^.

One of the basic characteristics that nonlinear interactions can induce in a system is multi-stability, commonly associated with the presence of a direct or indirect positive feedback loops in the network. Multi-stability is characterized by the dependence of the final steady state of the system on the initial conditions, and it has been observed experimentally *in vitro^20,21^, in vivo^22,23^*, and in synthetic circuits^24,25^. In the context of biological networks and drug treatment, this dependence on initial conditions may result in differences in the effect of a given drug, depending on the initial state of the system.

Here, we investigate whether the efficiency of drug inhibition is affected by the initial conditions in the proteins of signaling pathways. To do that, we set a computational high-throughput screening to explore all possible networks of 3-nodes and monitor their response to inhibition of one of the nodes. Each network has a topology that is represented by a system of three ODEs that represent a particular set of Michaelis-Menten interactions between input, target and output nodes. Each interaction is modeled as a Michaelis-Menten type of inteaction Starting from two different initial conditions, we generate two dose-response curves for each set of parameter values. The comparison of these two curves allows us to characterize each network topology in terms of its impact in the outcome of drug inhibition. Using this approach we found that, in most of the possible networks topologies, the initial state of the system determines the efficiency of a given drug, increasing, decreasing or even disrupting the efficiency of inhibition. We conclude that this dependence on the initial conditions may be compromising the reproducibility of *in vitro* and in vivo studies that involve inhibitory treatments.

## Results

### The strength of inhibition depends on the initial conditions for most of the networks

At first inspection, our screening reports differences between the two dose-response curves for around 80% of all network topologies. This suggests that the efficiency of the inhibition depends on the initial conditions for most of the possible three-node network topologies, at lest in certain region of the parameter space. The percentage of networks where the two dose-response curves do not coincide increases with the connectivity of the network, as shown in Fig. 1 (blue bars and left vertical axis), up to 97% for networks with 8 links between input, target and output (251 of all possible 256 networks of 8 links in our study). The percentage of simulations that show multiple dose-response curves also increases with the number of links in the network (green bars and right vertical axis in Fig. 1) up to 5.5% for the more connected topologies.

**Figure 1.**
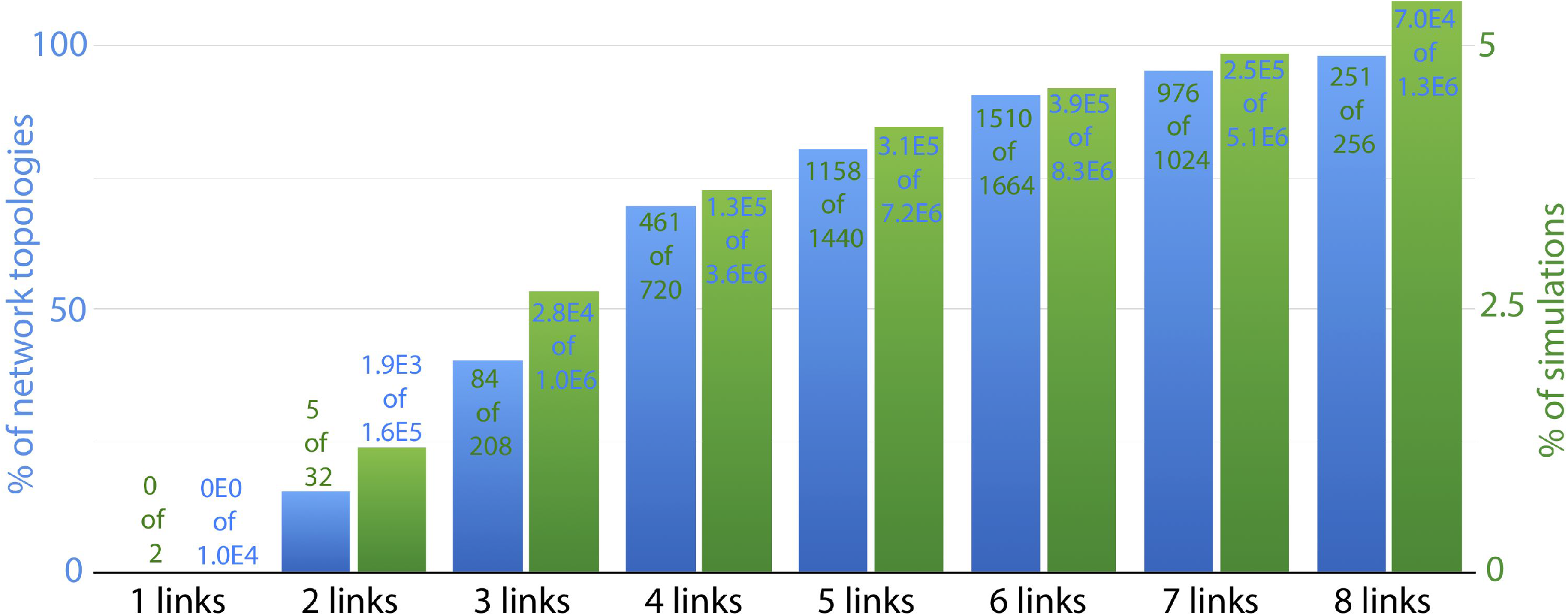
General statistical analysis of the high-throughput screening. Bar plot showing the percentage of cases with multiple dose-response curves to inhibition increases with the network connectivity. Blue bars correspond to the percentage of network topologies (left vertical axis) and green bars correspond to the percentage of simulations (right vertical axis) that show multiple dose-response curves (each simulation corresponds to a particular combination of parameters). Values in each bar illustrate the number of positive cases over the total number of cases.

When comparing the two dose-response curves, we can identify different scenarios of how the initial conditions affect the efficiency of the drug treatment. The most common scenario corresponds to a shift in the dose-response curve, i.e., the initial condition affects the efficiency of the inhibitor. This behavior is characterized by a shift in the *EC50* of the dose-response curve (i.e., the concentration of inhibitor that induces a half-maximal effect in the output). An example of this type of response is illustrated in Fig. 2a-d. The two dose-response curves are plotted in Fig. 2b, corresponding to each initial condition *IC_low_* and *IC_high_*, in blue (*D_Slow_*) and red (*DS_high_*), respectively. For this network configuration and these conditions, the EC50 of the inhibitor changes around 1.5 orders of magnitude. This type of dependence on the initial conditions is simply a result of a bistable regime, as shown in the phase plane in Fig. 2c (i.e., outside the bistable region, the final steady state does not depend on the initial condition whereas inside bistable regions different initial conditions may lead to different steady states). Inside the bistable regime, the nullclines for the inhibitor concentration marked in Fig. 2b show two stable fixed points coexisting for the same conditions (blue and red solid circles) and the unstable fixed point (empty black circle). Fig. 2d shows the bifurcation diagram with two stable branches that coexist for a particular range of values of inhibitor. Video S1 is an animation of how the nullclines and the steady states change with the concentration of inhibitor (black curves plot the trajectories of the initial conditions towards their corresponding steady state). This scenario can also occur in conditions where the inhibitor is acting as an activator of the output node, as illustrated in Supp. Fig. 4b.

**Figure 2.**
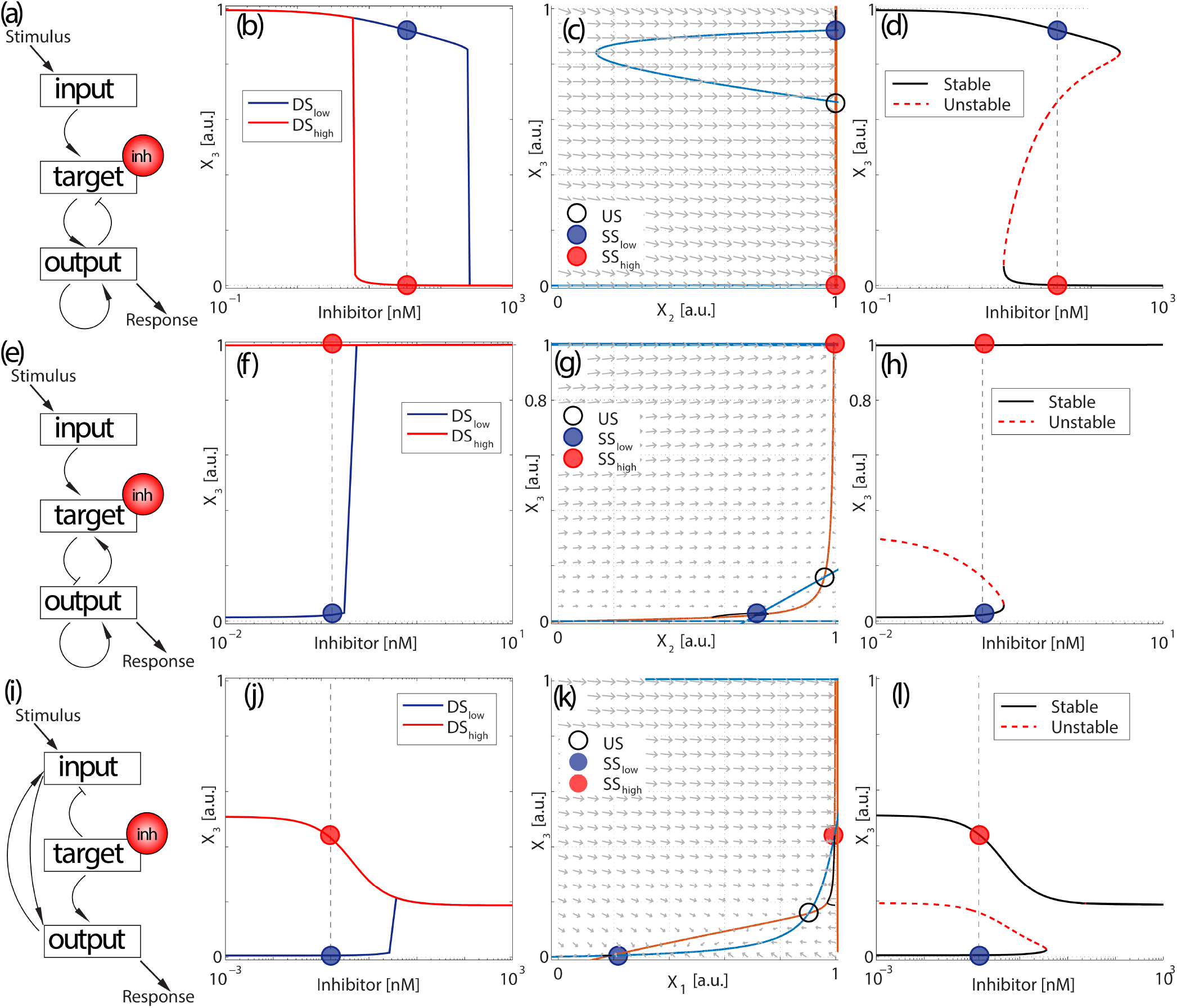
The effect of initial conditions can shape the dose-response curve in different ways. (a-d) Shift in the EC50, (e-h) insensitization of one of the dose-response curves, (i-l) Switch in the effect of the drug. Panels (a,e,i) represent the network topologies for each mode. Pointed arrows represent positive interactions (activation) and blunt arrows represent negative interactions (de-activation). Panels (b,f,j) represent the dose-response curves *DS_low_* (blue) and *DS_high_* (red) for initial conditions *IC_low_* and *IC_high_*, respectively. The rest of parameter values are the same between the two curves. Circles represent the steady state solutions of the system (blue and red solid circles correspond to *SS_low_* and *SS_high_*, respectively, and the empty black circle represents the unstable steady state) for a particular concentration of inhibitor (indicated by the vertical dashed lines in panels b,f,j). Panels (c,g,k) show the phase plane with vector field and nullclines for *X*_3_ (c,g,k) in blue and *X*_2_ (c,g) or X_1_ (k) in orange, representing the two stable steady states *SS_low_* (blue) and *SS_high_* (red) respectively. Panels (d,h,l) show the bifurcation diagram of *X*_3_. Black curves are the stable branches and the dash red curve is the unstable branch.

Another common scenario corresponds to one of the dose-response curves showing a standard response to treatment, while the other is not responding for the same range of concentrations of inhibitor. An example of these dual two dose-response curves is shown in Fig. 2f for the network illustrated in Fig. 2e. In this scenario, the inhibitor acts as an activator of *X*_3_ when we start from *IC_low_*, but if the system starts from *IC_high_*, it remains insensitive to changes in the concentration of inhibitor.

Alternatively, different initial conditions can also reverse the effect of a given drug. For instance, the same treatment can result in inhibition or activation of the output signal, simply depending on the initial state of activation of input, target and output nodes. An example of this behavior is shown in Fig. 2i-l. The two dose-response curves in Fig. 2j for the network in Fig. 2i show one of the curves (*DS_low_*) increasing when we increase the concentration of inhibitor, while the other (*DS_high_*) decreases. The phase diagram (Fig.2k) for intermediate values of the inhibitor shows two stable fixed points (filled red and blue circles), and an unstable fixed point (empty black circle). The bifurcation diagram (Fig.2l) presents two stable branches, with the upper branch decreasing when the inhibitor is increased. This diagram shows that the increase in *DS_low_* is caused by a transition from a bistable to a monostable regime with higher *X*_3_. This discontinuous jump in the dose-response curve is less pronounced for networks with higher connectivity, but we selected this example since its simplicity allows us to illustrate its nullclines in a two-dimensional phase plane, instead of a three-dimensional plot.

### Different initial conditions can induce increased or decreased treatment efficiency

Among all motifs that induce multiple dose-response curves, we can further characterize the topologies in terms of the comparison between the two curves with respect to the two initial conditions. The most common scenario corresponds to the situation illustrated in Fig. 2b and Video S1, where the less sensitive curve (higher *EC*50) corresponds to the initial condition that results from applying a low concentration of inhibitor *IC_low_*, and the more sensitive curve occurs when the system starts from the initial condition that results from applying a high concentration of inhibitor *IC_high_*. This increased sensitivity at intermediate concentrations of inhibitor occurs whether the treatment results in deactivation (as in Fig. 2b) or activation (as in Supp. Fig. 4b) of the target. This situation occurs because, in the bistable regime, each initial condition *IC_low_* and *IC_high_* evolves to its closest steady state in the phase space.

Several network topologies also exhibit a different scenario, characterized by an inversion in the sensitivity of the treatment between the two dose-response curves. This scenario is presented in Fig. 3b, and shows the red (*DS_high_*) and blue curves (*DS_low_*) swapped compared to Fig. 2b. For the inhibitor concentration indicated in Fig. 3b, the initial condition with lower *X*_3_ (red rhomb in Fig. 3c) evolves towards the steady state with higher *X_3_* (red solid circle). On the other hand, the initial condition with higher *X*_3_ (blue rhomb) evolves to the steady state with lower *X*_3_ (blue solid circle). This is clearly shown by the trajectories (black dotted curves) corresponding to two simulations with the same exact parameter values, but starting from the two different initial conditions (*IC_low_* and *IC_high_*).

**Figure 3.**
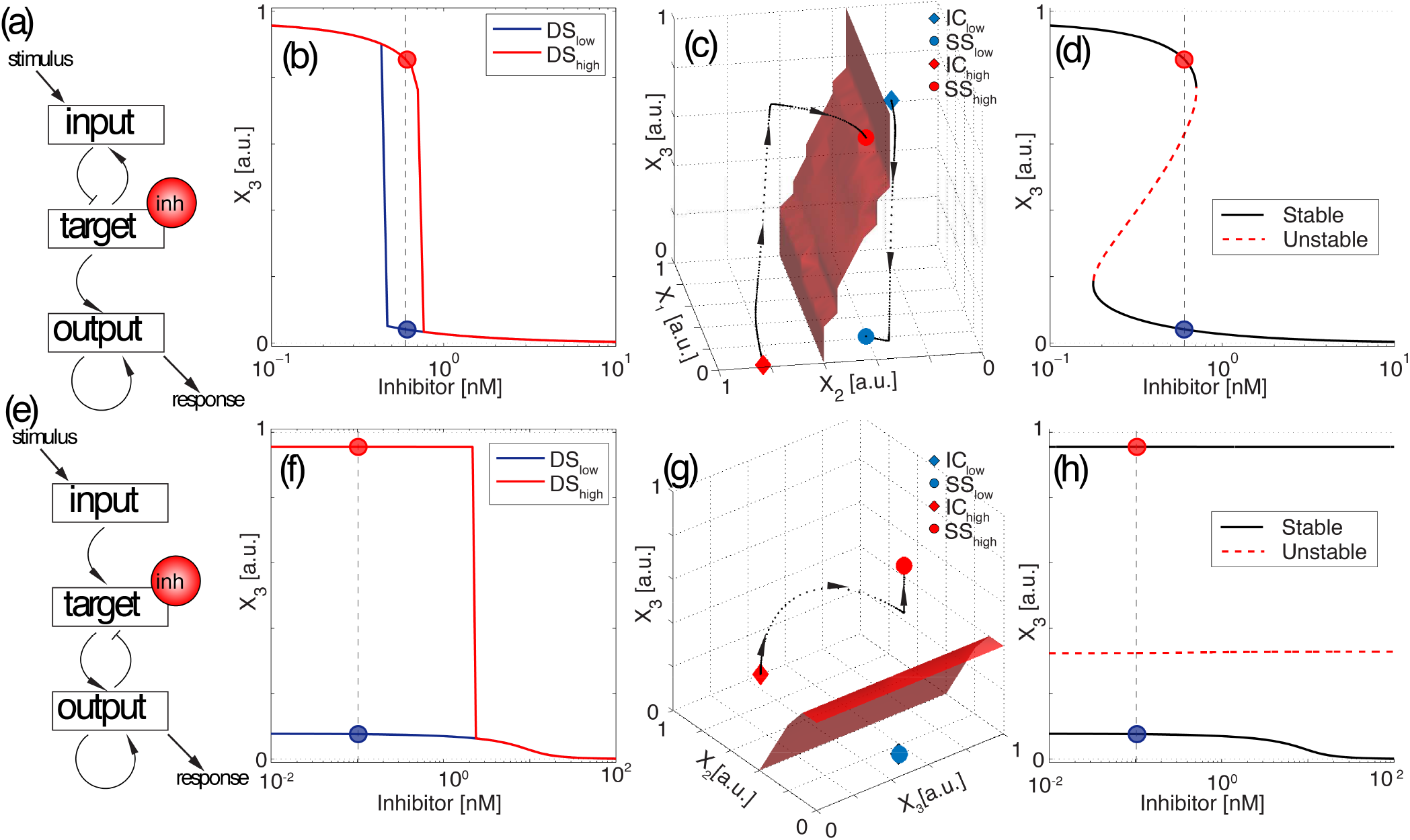
The network architecture can induce inverse bistability. (a-d) Shift in the EC50. (e-h) Insensitization of one of the dose-response curves. Panels (a,e) represent examples of network topologies that show two different cases of inverse bistability. Pointed arrows represent positive interactions (activation) and blunt arrows represents negative interactions (de-activation). Panels (b,f) represent the dose-response curves *DS_low_* (blue) and *DS_high_* (red) for initial conditions *IC_low_* and *IC_high_*, respectively. The rest of parameter values are the same for the two curves. Blue and red solid circles *SS_low_* and *SS_high_* represent the steady state solutions for a given concentration of inhibitor (vertical dashed line). Panels (c,g) represent the three-dimensional phase plane, with the trajectories of each simulation starting from each of the two initial conditions (red and blue rhombs), and the separatrix between the two basins of attraction (red surface). Panels (d,h) show the bifurcation diagram of *X*_3_. Black curves are the stable branches and the red dash curve is the unstable branch.

In terms of the effect of drug, the initial condition that results from applying a high concentration of inhibitor (*IC_high_*) shows a reduced response to the drug, compared to to the initial condition that results from a low concentration of inhibitor (*IC_low_*). In other words, the *EC*50 of *DS_high_* is now higher than *DS_low_*, as shown in Fig. 3b. This contrasts with the scenario of Fig. 2b and Video S1, where the EC50 of the drug is lower for *DS_high_* compared to *DS_low_*. To understand this behavior, we plot the three-dimensional separatrix between the two basins of attraction of the bistable regime in Fig. 3c. Since the separatrix divides the phase space vertically, the system is forced to perform a long path in *X*_3_ concentration towards the steady state in its basin of attraction. This is translated into a shift in the dose-response curves in the bistable regime, and therefore, an increase in the *EC*50 when the system is initially inhibited.

Since now each initial condition *IC_low_* and *IC_high_* does not transit to its closest steady state, but instead it evolves to the steady state that is further away in *X*_3_ concentration, we will refer to this scenario as inverse bistability. Video S2 is an animation of how the two initial conditions transit to their corresponding steady state for increasing concentrations of the inhibitor. This inversion of the bistable solutions, can also occur in conditions where the inhibitor is acting as an activator of the output node, as illustrated in Supp. Fig. 5b.

Analog to the situation of Fig. 2e-h where the dose-response curve (*DS_high_*) becomes insensitive to the drug, other topologies present the opposite scenario, i.e., the *DS_high_* responds to the drug but the *DS_low_* is insensitive. This scenario is illustrated in Fig. 3f for the network topology of Fig. 3e. Here, *DS_low_* responds by reducing *X*_3_ activation in less than 10%, while now a high initial concentration of inhibitor sensitizes the system, i.e., the dose-response curve (*DS_high_*) shows a much stronger inhibition of the output. Fig. 3g plots the three-dimensional phase space for a particular inhibitor concentration in the bistable regime. Again, the separatrix divides the space in such a way that the initial *IC_high_* evolves to the steady state with higher *X*_3_ and the switch in *DS_high_* (red line in Fig. 3f). Fig. 3h shows that the two branches are stable for all concentrations of inhibitor tested. Despite of this, the system is able to switch from one solution to the other because the separatrix moves relatively to the fixed initial conditions. Video S3 corresponds to an animation of this scenario. Please note that, depending on the parameter values, the same topology can exhibit different responses (for instance, the same network is used for Fig. 2a and Fig. 3e to generate normal and inverse irreversible bistability). The transition between different regimes depending on the parameter values is analyzed the next sections.

### The network architecture can induce inverse hysteresis loops

As discussed above, inverse bistability occurs due to the interplay in phase space between the initial conditions and the basins of attraction of the two final stable steady states. Nonetheless, our screening revealed another family of topologies that show an equivalent scenario of inverse bistability, but with additional features. An illustrative example of this behavior is shown in Fig. 4. The first example corresponds to a network topology of four links that shows inverse bistability as defined in the previous section, i.e, two dose-response curves where the *DS_high_* has a higher EC50 than *DS_low_*. Since *X*_2_ does not receive input from *X*_1_ and *X*_3_, the phase space is plotted in two dimensions to show the nullclines and the vector field (Fig. 4c). Interestingly, the bifurcation diagram in Fig. 4d shows a more complex configuration than in Fig. 3d, with the two stable branches now extending from low to high *X*_3_. This configuration induces another interesting property to these types of networks: Inverse bistability does not only occur when we start with fixed initial conditions, but also if the concentration of the inhibitor is gradually increased or decreased from each initial condition. In other words, if the concentration of inhibitor is progressively increased or decreased, the system follows a hysteresis loop that is reversed compared to the standard hysteresis observed in magnetism, optical and other physical systems. To illustrate that, we developed an animation where the concentration of inhibitor is gradually increased and then decreased, and the evolution of steady states form an inverse hysteresis loop (Video S4).

**Figure 4.**
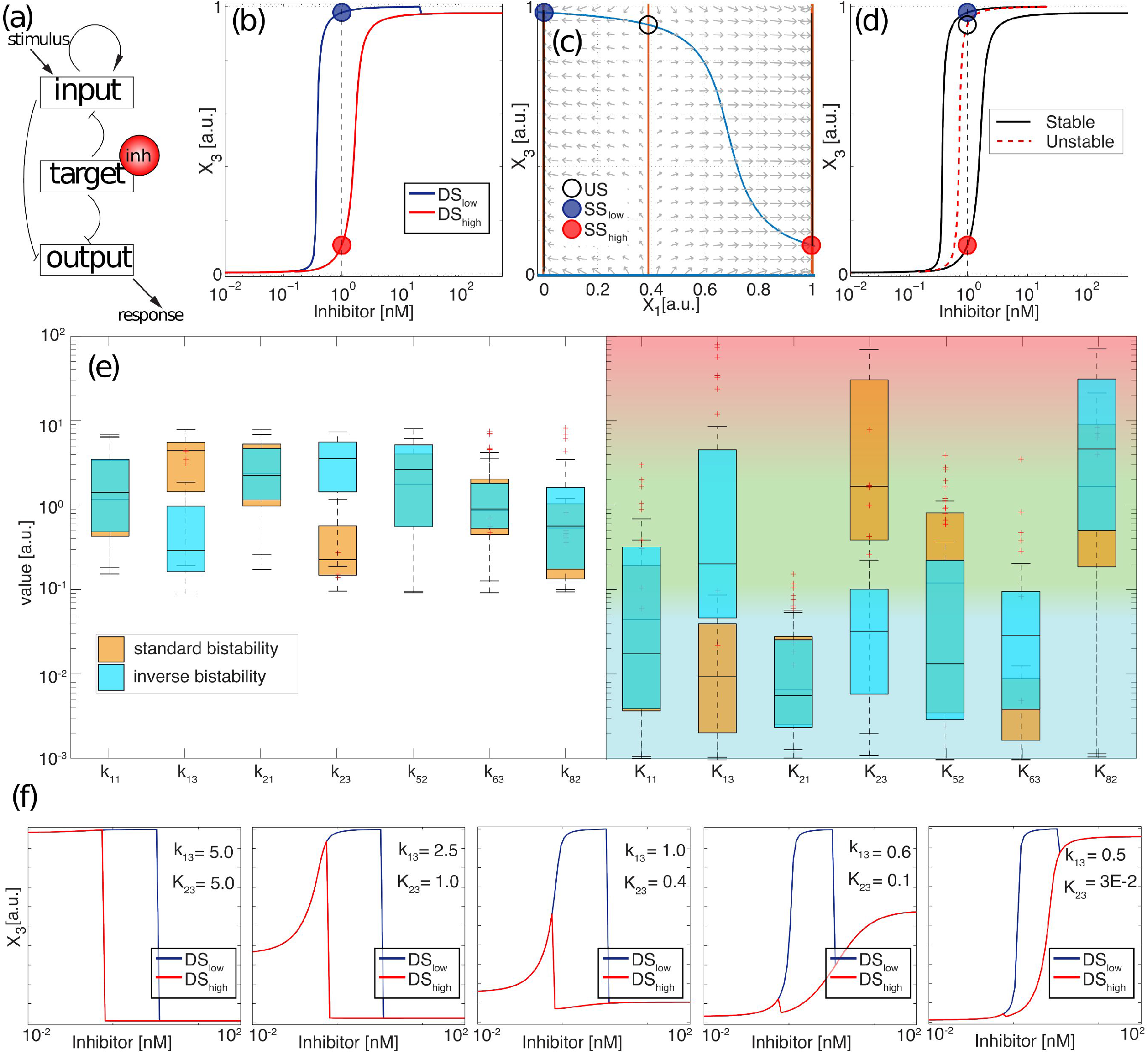
The network architecture can induce inverse hysteresis. (a) Example of a network architecture that induces inverse hysteresis. Pointed arrows represent positive interactions (activation) and blunt arrows represents negative interactions (de-activation). (b) Dose-response curves *DS_low_* (blue) and *DS_high_* red for initial conditions *IC_low_* and *IC_high_*, respectively. The rest of parameter values are the same between the two curves. Blue and red solid circles represent the two steady state solutions for a given concentration of inhibitor *(SS_low_* and *SS_high_*). (c) Phase plane with vector field and nullclines for *X*_3_ (blue) and *X*_1_ (orange). The black empty circle shows the unstable steady state. (d) Bifurcation diagram of *X*_3_. Black curves are the stable branches and the red dash curve is the unstable branch, for the inhibitor concentration indicated in panel b. (e) Box plot for all parameter sets that show standard and inverse hysteresis. Blue, green and red background represents the saturated, unconstrained and linear regimes of the Michaelis-Menten kinetics, respectively. (f) Changes in the dose-response curves when two parameters are varied from standard to inverse hysteresis conditions.

To understand the interactions that induce this inverse hysteresis response, we compared (Fig. 4e) 100 different sets of parameters in a box plot where this topology produces standard (orange) and inverse bistability (blue). This plot allows us to see that most values show overlapping distributions for both types of bistability, while two of them are clearly separated (*K*_2,3_ and *k*_1,3_ for this particular network). Next, dose-response curves are generated by changing these two parameters between their average values that produce standard or inverse bistability (the rest of parameters are fixed and correspond to the average of the mean for both orange and blue distributions). This analysis reveals that *K*_2,3_ mainly affects the response of *X*_3_ in the range of low inhibition, *k*_1,3_ mainly affects the steady state in the range of high inhibition, while the intermediate bistable region remains almost unchanged. When both are simultaneously varied (Fig. 4f), we clearly observe that these changes in low and high range of inhibitor interplay to change the nature of the drug from inhibitor to activator of the node *X*_3_.

This sequence also illustrates that, in some particular topologies, standard bistability can be converted to inverse bistability by manipulating some key interactions that reverse the effect of the inhibitor while maintaining the bistable region at intermediate inhibitor concentrations. To do that in this particular topology, the strength of the interaction between *X*_1_ and *X*_3_ is reduced, while the interaction between *X*_2_ and *X*_3_ goes from a linear to a unconstrained regime. A different topology with a similar transition from standard to inverse bistability is shown in Supp. Fig. 6. an additional example of a network topology able to produce inverse bistability and inverse hysteresis is shown in Supp. Fig. 8,

### Overall characterization of the topologies reveals the minimal motifs that exhibit inverse bistability

To characterize the basic ingredients underlying the inverse bistability, we proceed to analyze all potential topologies that exhibit this behavior and find relationships and similarities between them. When grouped by number of links, we observe that the percentage of networks that exhibit inverse bistability increases with the connectivity of the network (yellow columns in Fig. 5a); this also happens for the percentage of simulations (one simulation corresponds to one combination of parameters) showing inverse bistability (see Supp. Fig. 3). Fig. 5b represents all topologies that show inverse bistability as an atlas that correlates topologies by their architecture by identifying topologies that contain another topology of lower connectivity. This representation reveals 19 minimal motifs of 4 links that are contained in most of the higher connected topologies (the number of links in a given motif corresponds to the number of nonzero entries of the first 3 rows of the interaction matrix). These 19 topologies are represented as 3×3 matrix plots (which correspond to the three first rows of the interaction matrix) as follows: white is “1” (activation), black is “−1” (deactivation) and grey is “0” (no interaction)). The 4-link topologies that can also induce an inverse hysteresis loop are highlighted in red. The row below groups these 19 topologies in sets that only differ by two interactions (a given topology can result in different modes of response).

**Figure 5.**
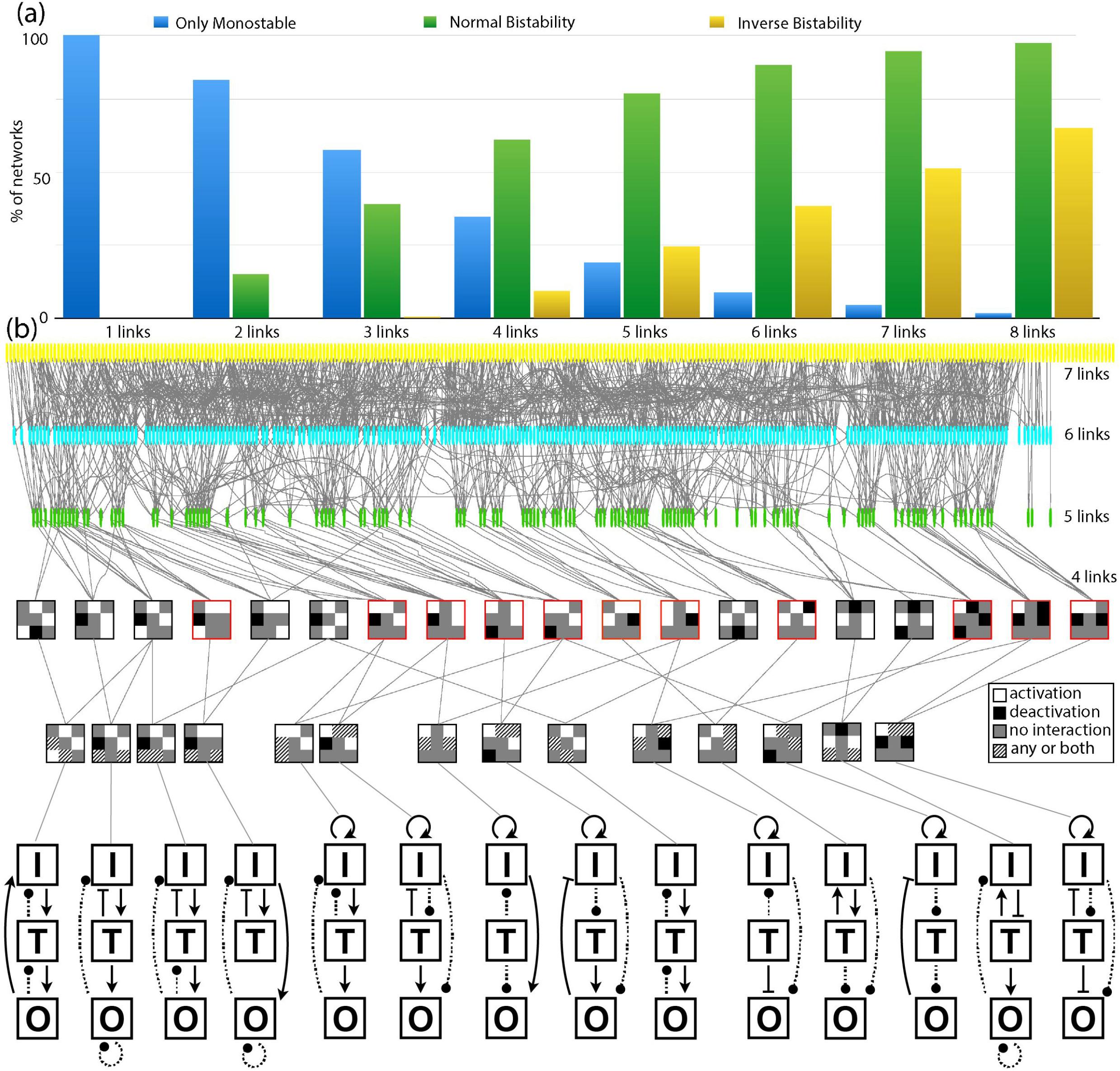
Characterization of networks that show inverse bistability and inverse hysteresis. (a) The percentage of topologies that show inverse bistability increases with the network connectivity. Blue bars correspond to the percentage of topologies with the same dose-response curve for both initial conditions; green and yellow bars correspond to the percentage of topologies that show increase or decrease of the EC50, respectively. (b) Atlas for all network topologies that induce inverse bistability. Circles represent each of the topologies where our screening has shown inverse bistability. Networks of different connectivity are represented in different colors. Gray lines link topologies that contain another topology of lower connectivity. Networks of lower connectivity are represented as matrixplots for the interactions, where white represents activation, black is deactivation and grey means no interaction. These minimal networks are then grouped in families where just one or two interactions change (marked with diagonal lines). Matrix plots highlighted in red correspond to topologies that can also produce inverse hysteresis loops. The topology corresponding to each matrixplot is shown below (interactions that vary in sign or in terms of presence/absence inside a family of minimal network topologies are represented with dashed lines).

All 19 minimal topologies combine positive and negative interactions (i.e, no networks where all interactions are positive or negative). In addition, all of them contain at least a positive feedback that can be direct or indirect (i.e, the self-activation of a node involves another node of the network). The negative interaction can take the form of an indirect feedback loop (as in Supp. Fig. 8a), an incoherent feed-forward loop, or not being part of a loop at all (as in Fig. 4a). We have found topologies where the interactions modulated by the inhibitor can either influence a positive, a negative feedback, a feedforwad loop, and even several of them simultaneously. We suggest that inverse bistability results from the interplay between the positive feedback (that generates the bistability) and the negative interactions that shape the basins of attraction. Additionally, the inhibitor has to directly or indirectly affect the positive feedback and induce a change between the two stable states at a given concentration.

## Discussion

In this paper, we present the first global analysis to study how the network topology influences drug treatment. To do that, we focus on small networks of three interacting nodes where one of the nodes is the target of a small molecule inhibitor. We compare dose-response curves of the same treatment starting from two different initial conditions. Our analysis reveals that the initial conditions affect the efficiency of the treatment in most network topologies of three nodes. This dependence arises from the nonlinear characteristics of the network topology, and it is translated into modifications in the dose-response curves and changes in the *EC*50 as well as in the overall effect of the inhibitor. Moreover, we found network configurations that show a novel behavior characterized by the inversion of the steady states respect to the initial conditions. In some conditions, this “inverse bistability” can also result in “inverse hysteresis loops”, where the reduction of the efficiency of the treatment also occurs when the concentration of inhibitor is varied gradually. To our knowledge, this is the first evidence of these type of responses. Finally, our study shows that most of the topologies that show this inverse bistability and hysteresis contain core motifs of four links composed by a positive feedback and a negative regulation.

The workflow of our high-throughput screening is an *in silico* simulation of the experimental workflow used to determine dose-response curves. The fact that all the points in a dose-response curve start from the same initial state interplays with the bistable regions generated by a given network topology, resulting in a complex scenario where the relation between the initial states and the basins of attraction in the phase space induces reversible or irreversible inverse bistability, and even inverse hysteresis loops.

When comparing the dose-response curves in standard versus inverse bistability, *DS_high_* has a higher sensitivity (reduced EC50) than *DS_low_* in standard bistability, while *DS_high_* has a lower sensitivity (increased EC50) than *DS_low_* in conditions of inverse bistability. This reduction in sensitivity is very different from the well-studied homologous or heterologous desensitization after repeated or prolonged receptor stimulation^29,30^. Receptor desensitization is achieved mainly by a single negative feedback loop that reduces the number or the efficiency of receptors on the cell surface after a initial stimulation^31,32^. This is translated into a initial strong transient activation of the targets downstream, while a second application of the stimulus does not show the same transient activation. While receptor desensitization focuses on transient responses, the reduced sensitivity resulting from the inverse bistability refers to true final steady states of the network.

Our study is limited to topologies of three main nodes that play different roles in the network, in an attempt to identify the minimal motifs that induce these dual dose-response curves. In principle, our results also apply to more larger networks with increasing number of nodes that interact linearly, since linear protein-protein interactions can be reduced to smaller networks with equivalent dynamics without reducing the spectrum of reported behaviors^3,33,34^. We also expect that larger more complex biological networks that contain any of the smaller motifs reported in our analysis (Fig. 5) will exhibit a similar or even stronger dependence on initial conditions (see for instance^35^), since our analysis shows that the percentage of the networks with multiple dose-response curve increases with the connectivity of the network. Our lab is currently working in the experimental validation and characterization of signaling pathways with dual response-curves when targeted by an inhibitor (manuscript in preparation)

The characterization of the effect of a drug starts with an accurate and reproducible *in vitro* or *in vivo* dose-response curve to establish the optimal dose or the optimal schedule or treatment. The fact that, for most topologies, different initial conditions give different dose-response curves may compromise the reproducibility of drug treatment between biological samples or even patients. In conclusion, when designing drugs and treatments that target proteins embedded in highly inter-connected networks such as signal regulatory pathways, the efficiency of a given compound cannot be predicted if the state of activation of the network is unknown.

## Methods

To study all potential network topologies that induce multiple dose-response curves, we set up a high-throughput approach that explores all possible network topologies, or connections between an input, a target and an output node, including positive and negative feedback auto-regulation (see Fig. 6a). This computational screening is inspired by previous studies that focus on network topologies inducing adaptation^26^, bistability and ultrasensitivity^27^ and spatial pattern formation^28^. Our approach introduces the effect of a drug inhibitor in one of the nodes of the network, and focuses mainly on the characterization of the effect of the network in shaping the response to the inhibition. To do that, we generate and compare dose-response curves for a given topology and set of parameters, but starting from different initial conditions.

**Figure 6.**
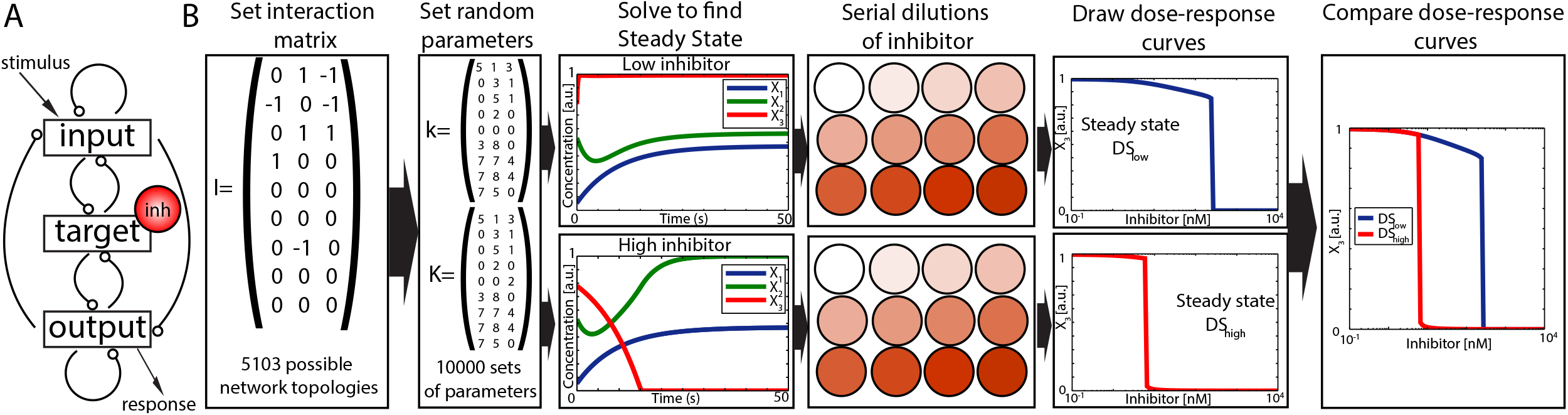
Scheme of the workflow for the high-throughput screening. (A) Scheme of the core network with all possible interactions between input, target, and output. (B) For each possible interaction matrix (5103 possible topologies) the phase space is sampled by randomly generating 10000 sets of parameter values for the catalytic (*k*), Michaelis-Menten (*K*) matrices. For each of these parameter sets, the three differential equations for input, target and output are solved numerically for two different inhibitor concentrations ([*inh*]_*low*_ = 0 nM and [*inh*]_*high*_ = 10^3^ nM). The resulting steady states are used as initial conditions (*IC_low_* and *IC_high_*) for numerical simulations applying a range of inhibitor concentrations. The steady state value of the output node (*X*_3_) is plotted against the inhibitor concentration to generate dose-response curves *DS_low_* and *DS_high_*. Finally, both dose-response curves are compared.

Our core network is composed of three main components: an input node that receives a constant external stimulus, a target node that is inhibited by the drug, and the output node, which is used as a readout of the system activity. Details of the dynamics of the interaction between the nodes and automatization of the screening are described in the Supp. Material. In brief, the set of interactions can be generalized in the following equation:

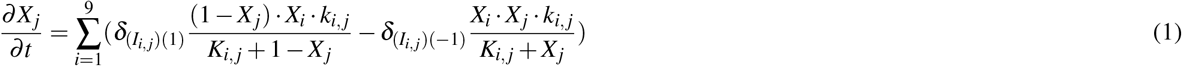

where *X* is the state vector that contains the concentration of the active version of the input *X*_1_, target *X*_2_ and output *X*_3_, as well as the value of the background activator (*X*_4_,*X*_5_,*X*_6_), and a deactivator (*X*_7_,*X*_8_,*X*_9_) enzymes for each of them. These background interactions are incorporated to ensure that each node receives at least one activating and one deactivating interaction (see Supp. Material). Therefore, independently of the number of links, the system of equations for each topology only contains three equations (the concentrations *X*_4_,…,*X*_9_ are constants (parameters) rather than variables). *I_i,j_* represents the components of the interaction matrices for all 5103 possible networks of interactions between input, target and output in our study (see Supp. Material). Here, a given component *I_i,j_* of the matrix is zero if *X_i_* does not affect *X_j_*, 1 if the *X_i_* activates *X_j_* and −1 if *X_i_* deactivates *X_j_*. *δ*_(*I_i,j_*)(1)_ and *δ*(*I_i,j_*)(−1) are Kronecker delta functions that are 1 when the value *I_i,j_* is 1 or −1, respectively. This way, the left part of the sum is nonzero when the component *X_i_* activates *X_j_*, while the right side is nonzero when *X_i_* deactivates *X_j_*.

Parameters *k,_i,j_* and *K_i,j_* correspond to the catalytic and Michaelis-Menten constants for the activation or deactivation of *X_j_* by *X_i_*. The effect of inhibitor is incorporated as a sigmoidal function, assuming fast dynamics of binding and unbinding to its target (quasi-steady state approximation) and that the inhibitor is in excess over the enzyme *X*_2_ (see Supp. Material). Considering this, *X*_2_ is substituted by the expression

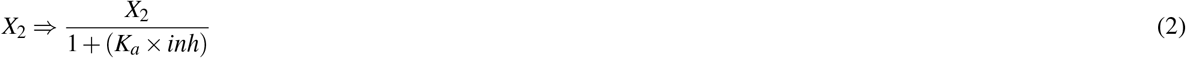

when *i* = 2 in Eq. 1 (i.e., whenever *X*_2_ acts as an activating or deactivating enzyme). This way, the effect of the inhibitor can be directly incorporated into the equations independently of the architecture of the network, strongly facilitating the screening process. We fix the value of *K_a_* of our inhibitor as 10^7^ 1/*M* (see Supp. Material), and assume that it is in excess over the enzyme *X*_2_. This approach excludes from our screening all topologies where *X*_2_ regulates itself (see Supp. Material).

The workflow can be described as follows (see Fig. 6b): For each particular network topology, all components *k_i,j_* and *K_i,j_* of the catalytic and Michaelis-Menten constant matrices are randomly set from a desired range of values (see Supp. Material). Then, the system is numerically solved for two different constant concentrations of inhibitor ([*inh*]_*low*_ = 0 nM and [*inh*]_*high*_ = 10^3^ nM), and the resulting steady state values of input, target and output are then used as initial conditions for new numerical simulations. Next, different concentrations of inhibitor are applied to the networks starting from these two initial conditions. The steady state of the output *X*_3_ is used to draw the corresponding dose-response curve. Finally, the two dose-response curves are analyzed, compared and classified (see Supp. Material for a more detailed explanation). This is repeated 10000 times for each topology, with different catalytic and Michaelis-Menten constant matrices, to sample the parameters space and identify regions where the dose-response curve depends on the initial conditions. Based on this, all network topologies are classified depending on the relationship between the two dose-response curves. This way, if both curves *DS_low_* and *DS_high_* are identical, the response of the inhibitor does not depend on the initial conditions, while if the two curves are different, this means that the effect of the inhibitor is dependent on the initial state of activation of the system.

This workflow is designed to mimic the typical experimental methodology to determine dose-response curves: Starting from multiple equivalent samples in the same experimental condition, different concentrations of the drug are administered to each of the samples, and the final steady state of the readout is plotted against the concentration of the drug. This is different from the typical studies of bistability in physical and chemical systems^20,21^, where an input parameter is gradually increased or decreased (i.e., the initial condition for each point in the curve corresponds to the steady state of the previous point in the analysis).

## Acknowledgements (not compulsory)

This work has been supported by the Ministry of Science and Technology of Spain via a Ramon y Cajal Fellowship (Ref. RYC-2010-07450), a grant from Plan Nacional framework (Ref. BFU2011-30303 and & BFU2014-53299-P) and a FPU fellowship. We thank Raul Guantes, Juan Diaz Colunga, Marta Ibanes, Rosa Martinez Corral, Saul Ares and Katherine Gonzales for invaluable help and technical assistance.

## Author contributions statement

DGM and VDM: Designed research, performed research, wrote the manuscript. All authors reviewed the manuscript.

